# Differential cellular immune responses against *Orientia tsutsugamushi* Karp and Gilliam strains following acute infection in mice

**DOI:** 10.1101/2023.06.07.543994

**Authors:** Joseph D Thiriot, Yuejin Liang, Casey Gonzales, Jiaren Sun, Lynn Soong

## Abstract

Scrub typhus is the leading source of febrile illness in endemic countries due to infection with *Orientia tsutsugamushi* (*Ot*), a seriously understudied intracellular bacterium. Pulmonary complications in patients are common and can develop into life threatening conditions. The diverse antigenicity of *Ot* genotypes and inter-strain differences seem to be connected to varied virulence and clinical outcomes; however, detailed studies of strain-related pulmonary immune responses in human patients or experimental animals are lacking. In this study, we used two clinically prevalent bacterial strains, Karp and Gilliam, and revealed cellular immune responses in inflamed lungs and potential biomarkers of disease severity. We found that outbred CD-1 mice were highly susceptible to both Karp and Gilliam strains; however, C57BL/6 (B6) mice were susceptible to Karp, but resistant to Gilliam (with self-limiting infection), corresponding to their tissue bacterial burdens and lung pathological changes. Multicolor flow cytometric analyses of perfused B6 mouse lungs revealed robust and sustained influx and activation of innate immune cells (monocytes, macrophages, neutrophils, and NK cells), followed by those of CD4^+^ and CD8^+^ T cells, during Karp infection, but such responses were greatly attenuated during Gilliam infection. The robust cellular responses in Karp-infected B6 mice were positively correlated with significantly early and high levels of serum cytokine/chemokine protein levels (CXCL1, CCL2/3/5, and G-CSF), as well as pulmonary gene expression (*CXCL1/2, CCL2/3/4,* and *IFNγ*). *In vitro* infection of B6 mouse-derived primary macrophages also revealed bacterial strain-dependent immune gene expression profiles. This study provided the first lines of evidence that highlighted differential tissue cellular responses against Karp vs. Gilliam infection, offering a framework for future investigation of *Ot* strain-related mechanisms of disease pathogenesis vs. infection control.

**Authors Summary:** *Orientia tsutsugamushi* (*Ot*) infection-induced scrub typhus is a leading cause of febrile illness in endemic countries. Research on *Ot* strain-related disease outcomes or immune signatures in tissue and blood samples is very limited. Using two clinically prevalent strains (Karp and Gilliam), we examined host susceptibility in inbred and outbred mouse models and provided new evidence for the activation of pulmonary immune cell subsets during the acute stages of infection. While Gilliam-infected C57BL/6 (B6) mice developed self-limiting infection, mild cellular responses, and tissue injury, Karp infection led to a strong and sustained activation of innate immune cells, followed by extensive influx of activated T cells, which correlated to protein levels of inflammatory cytokines/chemokines in serum samples. We also provided *in vitro* evidence for *Ot* strain-dependent immune gene profiles, indicating differential macrophage responses to Karp versus Gilliam bacteria. This is the first comparison of different scrub typhus mouse models with in-depth analyses of cellular responses in inflamed lungs, offering novel insights into potential mechanisms of disease progression versus infection control related to *Ot* strains and laying the foundation for future investigations.

## Introduction

*Orientia tsutsugamushi* (*Ot*) is a severely neglected, tropical bacterial pathogen that causes the most prevalent febrile illness in the Rickettsiales order. It is estimated that this infection results in 150,000 global deaths per year and 1 million cases annually, encompassing over 1 billion people at risk of infection [1-4]. This trombiculid mite-transmitted bacterium can cause subclinical infection or non-specific symptoms, including headache, fever, myalgia, rash, and regional lymphadenopathy [5]. Patients with delayed or inappropriate treatment can develop into lethal cases that lead to acute respiratory distress syndrome, acute encephalitis syndrome, and multiorgan failure [3, 5, 6]. There are no approved vaccines currently for scrub typhus, mainly due to the paucity of knowledge associated with bacterial pathogenesis and host immunity. Research efforts and resources to study human immune responses to *Ot* infection and tissue-specific changes during disease progression are very limited, generally confined to analyses of patient sera and peripheral blood cells [7-10]. Since the lungs of infected patients and experimental animals carry the highest bacterial burdens and pathological changes [11-14], there is a great need to reveal pulmonary innate and cellular immune responses, in the context of exposure to different *Ot* strains that are known to be associated with diverse disease outcomes.

Within the Rickettsiaceae family, *Ot* has a relatively large genome at 1.9-2.5 Mbp, with an astounding number of repeat sequences and conjugative elements [15]. This is in stark contrast to its closely related *Rickettsia* species that tend to have small (typically 1.1-1.3 Mbp) and surprisingly stable genomes [16]. This proclivity for repeating elements is not uncommon among other intracellular bacteria spp, such as *Wolbachia* or *Ehrlichia*, which possess mobile genetic elements and tandem intergenic repeats, respectively [17, 18]. The unique characteristics of *Ot* may account for its high diversity, and consequently antigenicity and virulence. Seven geographically diverse *Ot* genotype groups have been identified, each of them composed of multiple serotypes of strains, based on their major type-specific antigens (TSA56) [15, 19]. Multiple strains of *Ot* contribute to the overall global burdens of scrub typhus patients [15, 19]. While the geographic distribution of predominant strains is region specific, Karp and Gilliam strains are of particular interest, as they attribute for 65% and 26% of clinical cases in endemic countries, respectively and they can cause severe scrub typhus and death, even among antibiotic-treated patients [20]. Yet, the comparative study of host immune responses to these two strains is necessary to dissect the unique immune signature in scrub typhus.

Improved murine and non-human primate models of scrub typhus have been developed in recent years; some of these models closely resemble bacterial transmission, acute tissue injury, pathologic and immunologic features in patients [11, 12, 20-24]. Several murine-based reports conducted more than 10 years ago are especially relevant to this study, due to their consideration of antigenically distinct *Ot* strains in the context of *Ot* inoculation doses and/or host susceptibility. For example, while *Ot* TA716 strain led to non-lethal infection in BALB/c and C3H/He mice, a lower dose of Karp caused lethal infection in BALB/c and C3H/He mice; in contrast, a similar dose of Gilliam let to lethal infection in C3H/He mice, but not in BALB/c mice [25-27]. Yet, the mechanisms underlying such diverse infection outcomes have never been investigated at the tissue, cellular, or molecular levels. Given that Karp, Gilliam, and TA716 strains are all pathogenic in humans, causing severe clinical outcomes in some patients [28-30], a better understanding of the pathogenic mechanisms underlying host susceptibility to clinically prevalent *Ot* strains is in urgent need, especially in the context of advanced immunologic approaches and assays.

In this study, we investigated host susceptibility and clinical outcomes following inoculation with the comparable doses of Karp and Gilliam in C57BL/6 (B6) vs. outbred CD-1 mice. While CD-1 mice were highly susceptibility to both *Ot* strains, as we and others have reported [13, 14], B6 mice were susceptible to Karp, but highly resistant to Gilliam (with no signs of disease). Focusing on the B6 models, we perfused lungs and performed multi-color flow cytometric analyses of tissue-infiltrating immune cell subsets and their activation status. We revealed robust and sustained innate immune responses in Karp-infected mice, which positively correlated with cytokine levels in serum samples and the pulmonary proinflammatory gene expression profiles. Our data reveal *Ot* strain-dependent differences for lung innate and cellular immune responses in the context of distinct clinical outcomes of scrub typhus. This study contributes to the understanding of the cellular and molecular basis underlying immune responses and pathology related to the *Ot* strain, offering a framework for future investigations into disease control strategies.

## Materials and Methods

### Ethics Statement

The University of Texas Medical Branch (UTMB) complies with the USDA Animal Welfare Act (Public Law 89-544), the Health Research Extension Act of 1985 (Public Law 99-158), the Public Health Service Policy on Humane Care and Use of Laboratory Animals, and the NAS Guide for the Care and Use of Laboratory Animals (ISBN-13). UTMB is a registered Research Facility under the Animal Welfare Act. It complies with NIH policy and has current assurance with the Office of Laboratory Animal Welfare. All procedures were approved by the Institutional Biosafety Committee, in accordance with Guidelines for Biosafety in Microbiological and Biomedical Laboratories. Infections were performed following Institutional Animal Care and Use Committee approved protocols (2101001 and 1902006) at UTMB in Galveston, TX.

### Mouse infection and organ collection

Female Swiss Webster CD-1 outbred mice were purchased from Envigo (East Millstone, NJ). Female B6 mice were purchased from Jackson Laboratory (Bar Harbor, ME). Mice were maintained under specific pathogen-free conditions in the same room for 9-10 days and infected at 8-12 weeks of age. Infections were performed in the Galveston National Laboratory ABSL3 facility at UTMB. All tissue processing and analytic procedures were performed in BSL3 or BSL2 laboratories, respectively. All infections were performed using the same bacterial stock of *Ot* Karp or *Ot* Gilliam strain prepared from L929 cells, as described in our previous studies [31, 32]. Two independent studies were performed, in which B6 mice were inoculated i.v. with 5.6-6.8 × 10^4^ focus forming units (FFU) of Karp or Gilliam stocks (200 μl), or with PBS (negative controls). CD-1 mice were inoculated i.v. with 3.4 × 10^4^ FFU of the same Karp or Gilliam stocks (200 μl), or with PBS. Mice were monitored daily for body weight, signs of disease, and disease scores. The disease score (ranged from 0-5) was based on an approved animal sickness protocol [21, 32]. The criteria included mobility/lethargy, hunching, fur ruffling, bilateral conjunctivitis, and weight loss: 0-normal behavior; 1- active, some weight loss (<5%); 2- weight loss (6-10%), some ruffled fur (between shoulders); 3- weight loss (11-19%), pronounced ruffled fur, hunched posture, erythema, signs of reduced food/water taken; 4- weight loss (20-25%), decreased activity, bilateral conjunctivitis, showing signs of incapable to reaching food/water; 5- non-responsive (or weight loss of greater than 25%) animal need to be humanely euthanized. Blood was taken, lungs were perfused, and then lungs and spleen were collected at days 4, 8, 12, and 32; the corresponding lung lobes between samples were used for the below comparative studies. Samples (5/group) were inactivated for immediate and subsequent analysis, with mock samples serving as the controls.

### Bacterial load quantification

Animal tissues were collected and stored in RNA*Later* (Qiagen) at 4°C overnight for inactivation and then stored at −80°C. DNA was extracted from tissue or cell culture using the DNeasy Blood & Tissue Kit (Qiagen) following the manufacturer’s instructions. For tissues, less than or equal to 30 mg was used for each extraction. Bacterial burdens were quantified via qPCR and normalized to total nanogram (ng) of DNA per μL of sample [48]. The 47 kDa gene was amplified using the primer pair OtsuF630 (5′-AACTGATTTTATTCAAACTAATGCTGCT-3′) and OtsuR747 (5′-TATGCCTGAGTAAGATACGTGAATGGAATT-3′) primers (IDT, Coralville, IA). Data were expressed as the copy number of 47-kDa gene per ng of DNA. The copy number for the 47-kDa gene was determined by serial dilution of known concentrations of a control plasmid containing a single-copy insert of the gene.

### Lung immune cell flow cytometry analysis

After perfusion by using PBS, equivalent portions of lung lobes/tissues were harvested from infected and control mice, processed, and stained [21]. Briefly, tissues were minced and digested with 0.05% collagenase type IV (Gibco/Thermo Fisher Scientific) in Dulbecco’s Modified Eagle’s Medium (DMEM, Sigma-Aldrich, St. Louis, MO) for 30 min at 37°C. Minced tissues were homogenized via abrasion against cell strainer gauze. Lung single-cell suspensions were made by passing lung homogenates through 70-μm cell strainers. Red blood cells were removed by using Red Cell Lysis Buffer (Sigma-Aldrich). Leukocytes were stained with the Fixable Viability Dye (eFluor 506) (eBioscience/Thermo Fisher Scientific, Waltham, MA) for live/dead cell staining, blocked with FcγR blocker, and stained with fluorochrome-labeled antibodies (Abs). The following Abs purchased from Thermo Fisher Scientific and BioLegend (San Diego CA): PE-Cy7-anti-CD3ε (145-2C11), Pacific Blue-anti-CD4 (GK1.5), APC-Cy7-anti-CD8a (53– 6.7), APC-anti-Ly6G (1A8-Ly6G), PE-anti-CD80 (16-10A1), BV421-anti-CD206 (CO68C2), FITC-anti-CD64 (X54-5/7.1), PerCP-Cy5.5-anti-CD11b (M1/70), PE-anti-CD44 (IM7), FITC-anti-NK1.1 (PK136), PE-anti-CD63 (NVG-2) and FITC-anti-CD69 (H1.2F3). Cells were fixed in 2% paraformaldehyde overnight at 4°C. Data were collected by a BD LSR Fortessa (BD Bioscience, San Jose, CA) and analyzed using FlowJo software version 10 (Tree Star, Ashland, OR).

### Quantitative reverse transcriptase PCR (qRT-PCR)

Total RNA was extracted from tissues or cultured cells by using the RNeasy mini kit (Qiagen) and treated with DNase, according to the manufacturer’s protocol. cDNA was synthesized via the iScript cDNA synthesis kit (Bio-Rad). Target gene abundance was measured by qRT-PCR using a Bio-Rad CFX96 real-time PCR apparatus. SYBR Green Master mix (Bio-Rad) was used for all PCR reactions. The assay included: denaturing at 95°C for 3 min followed with 40 cycles of: 10s at 95°C and 30s at 60°C. The 2^−ΔΔCT^ method was used to calculate relative abundance of mRNA expression. Glyceraldehyde-3-phosphate dehydrogenase (GAPDH) was used as the housekeeping gene for all analyses. Primer sequences used are listed in **Table S1**.

### Histology

All tissues were fixed in 10% neutral buffered formalin and embedded in paraffin. Tissue sections (5 µm thickness) were stained with hematoxylin and eosin and mounted on slides, as in our previous reports [11, 23]. Sections were imaged under an Olympus BX53 microscope, and at least five random fields for each section were captured.

### Serum cytokine and chemokine levels

Whole blood was collected from B6 mice at days 4, 8,12, and 32 of infection and compared with mock controls. Serum was isolated and inactivated, as described in our previous study [23, 24]. The Pro Mouse Cytokine 23-Plex Kit (Bio-Rad) was used to measure cytokine and chemokine levels. The Bio-Rad Bio-Plex Plate Washer and Bio-Plex 200 machines were used for sample processing and analysis. All processes were completed following the manufacturer’s instructions.

### Infection of mouse bone marrow-derived macrophages (MΦ)

Bone marrow cells were collected from the tibia and femur of B6 mice and treated with red blood cell lysis buffer (Sigma Aldrich). MΦs were generated by incubating bone marrow cells at 37°C with 40 ng/ml M-CSF (Biolegend, San Diego, CA) in complete RPMI 1640 medium (Gibco) [21]. Cell medium was replenished at day 3, 6, and 9, and cells were collected at day 10. MΦs (5 × 10^5^) were seeded into 24-well plates and allowed to adhere for 3 h prior to infection. Bacteria were added at a multiplicity of infection (MOI) of 10 in 100 μl to the wells for 1 h with periodic rotating. Then, 900 μl of fresh media was added per well, and plates were incubated at 37°C with 5% CO_2_.

### Statistical Analyses

All *in vivo* data are representative of two independent experiments and are presented as mean ± SEM. Differences between control and infected groups were analyzed by using the Student’s t test and one-way ANOVA (parametric and non-parametric), where appropriate. Differences between survival curves were analyzed by using the Log-ranked (Mantel-Cox) test. GraphPad Prism software was used for data analysis. Statistically significant values are denoted as \**p* < 0.05, ** *p* < 0.01, *** *p* < 0.001, and **** *p* < 0.0001, respectively.

## Results

### Karp infection causes consistently higher disease severity and bacterial burdens in both B6 and CD-1 mouse models

To compare *Ot* strain-related virulence, we infected B6 mice with the same doses of Karp and Gilliam (6.8 ×10^4^ FFU, i.v.) and monitored the mice daily. As shown in **Fig 1**, Karp-infected mice began to lose body weight on day 4, with 10% of them succumbed to infection by day 12, while the survival mice stopped weight loss at day 13, indicating a sublethal outcome. In sharp contrast, mice infected with Gilliam did not exhibit any weight loss or disease scores, indicating a highly resistant phenotype.

**Fig 1.**
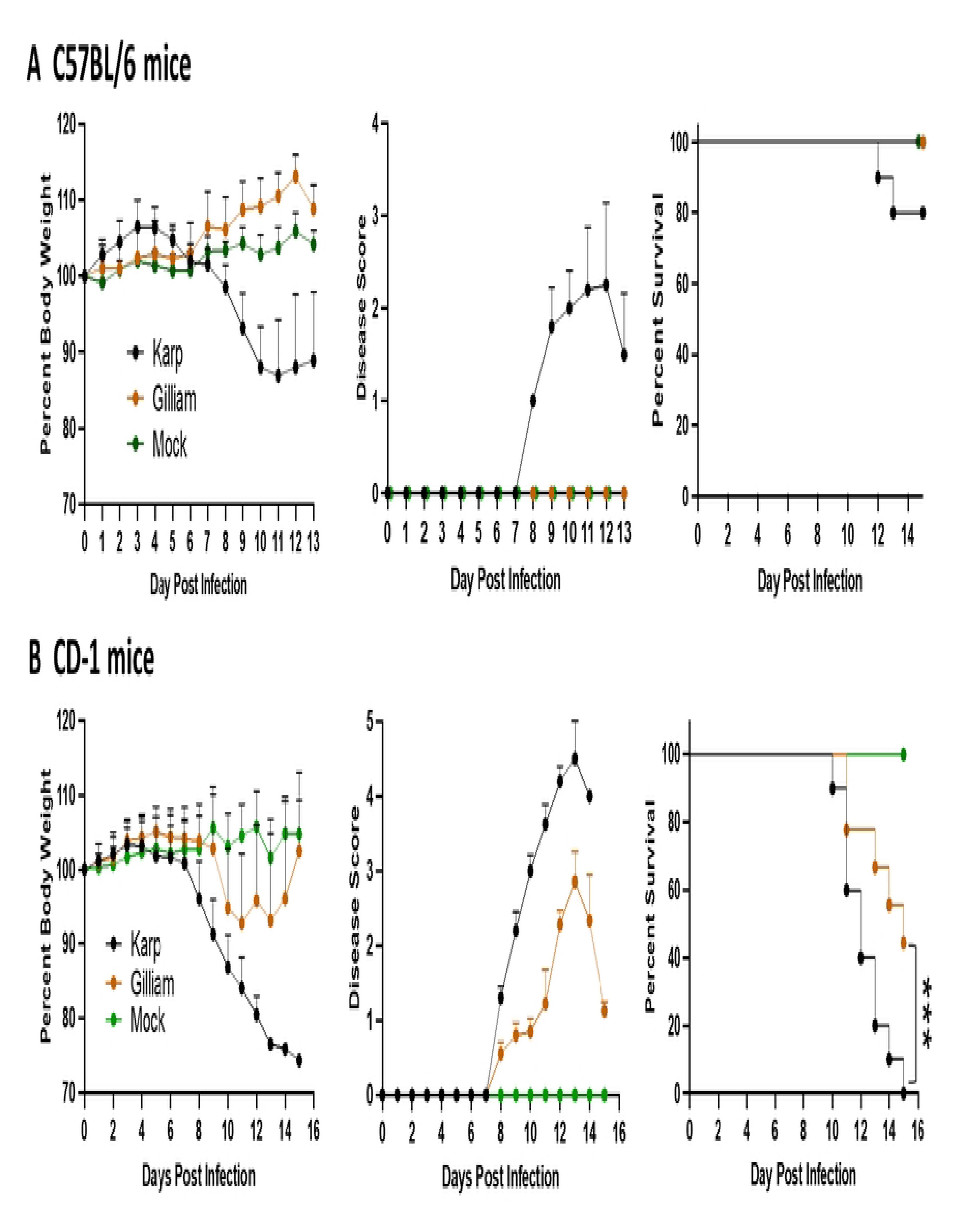
Diverse clinical outcomes following inoculation of *Ot* Karp and Gilliam in inbred versus outbred mouse models. A) C57BL/6 mice were inoculated with the Karp or Gilliam strain (5.6-6.8×10^4^ FFU, i.v., 10/group) or PBS (mock, 5/group). Mice were monitored daily for body weight changes (in percentage), clinical signs of disease score (see Materials & Method for detail), and percent of survival rate (%). B) CD-1 mice were infected similarly and side-by-side with those in A, but at a lower inoculation dose (3.4×10^4^ FFU, 10/group) or PBS (mock, 5/group). Mice were monitored daily, as in A. Shown are representative results from two independent studies with similar trends. ***, *p* < 0.001.

To validate the infectious nature of Gilliam strain and to explore differential host susceptibility to these two *Ot* strains, we also i.v. infected outbred CD-1 Swiss Webster mice with Karp or Gilliam strain (3.4 ×10^4^ FFU). This relatively low infectious dose was selected based on our recent publication, which has indicated higher susceptibility of CD-1 mice than B6 mice to Karp [14]. As shown in **Fig 1B**, CD-1 mice with Karp infection began to lose weight at day 8 and were completely moribund by day 15. CD-1 mice with Gilliam infection began to lose weight at day 10 and reached 50% mortality by day 15, with the remaining mice recovering gradually. These changes were consistent with the disease scores, showing that Karp-infected mice developed higher disease scores than Gilliam-infected ones. Collectively, the side-by-side comparison of inbred and outbred mouse models not only validated the infectivity of our bacterial stocks, but more importantly, indicated the diversity of bacterial virulence and host susceptibility in our established murine models (Karp-infected CD-1 > Gilliam-infected CD-1 >> Karp-infected B6 >> Gilliam-infected B6).

### Karp, but not Gilliam, infection causes severe pulmonary damage during acute infection

Having confirmed the diversity of host susceptibility to Karp vs. Gilliam infection, we decided to focus on the B6 models in the below study, for detailed analyses of host immune responses to two *Ot* strains. As shown in **Fig 2A**, bacterial burdens in Karp-infected lung tissues peaked at day 4, maintained at relatively high levels at day 8, and dropped substantially by day 12, with little to no burdens detected at day 32. In sharp contrast, bacterial burdens in Gilliam-infected lung tissues were relatively low (peaked at day 12). Karp-infected lung tissues possessed 928- and 221-fold higher bacterial burdens at days 4 and 8 than those of Gilliam-infected lungs (*p* < 0.0001), respectively. Very similar kinetic patterns were observed for Karp-and Gilliam-infected spleen samples; Karp-infected spleens had 9.69- and 8.32-fold higher bacterial burdens than those of Gilliam-infected spleens at day 4 and 8 respectively (**Fig 2B**). Therefore, Karp infection resulted in higher bacterial burdens in the lung and spleen tissues as compared to Gilliam infection during the acute stages.

**Fig 2.**
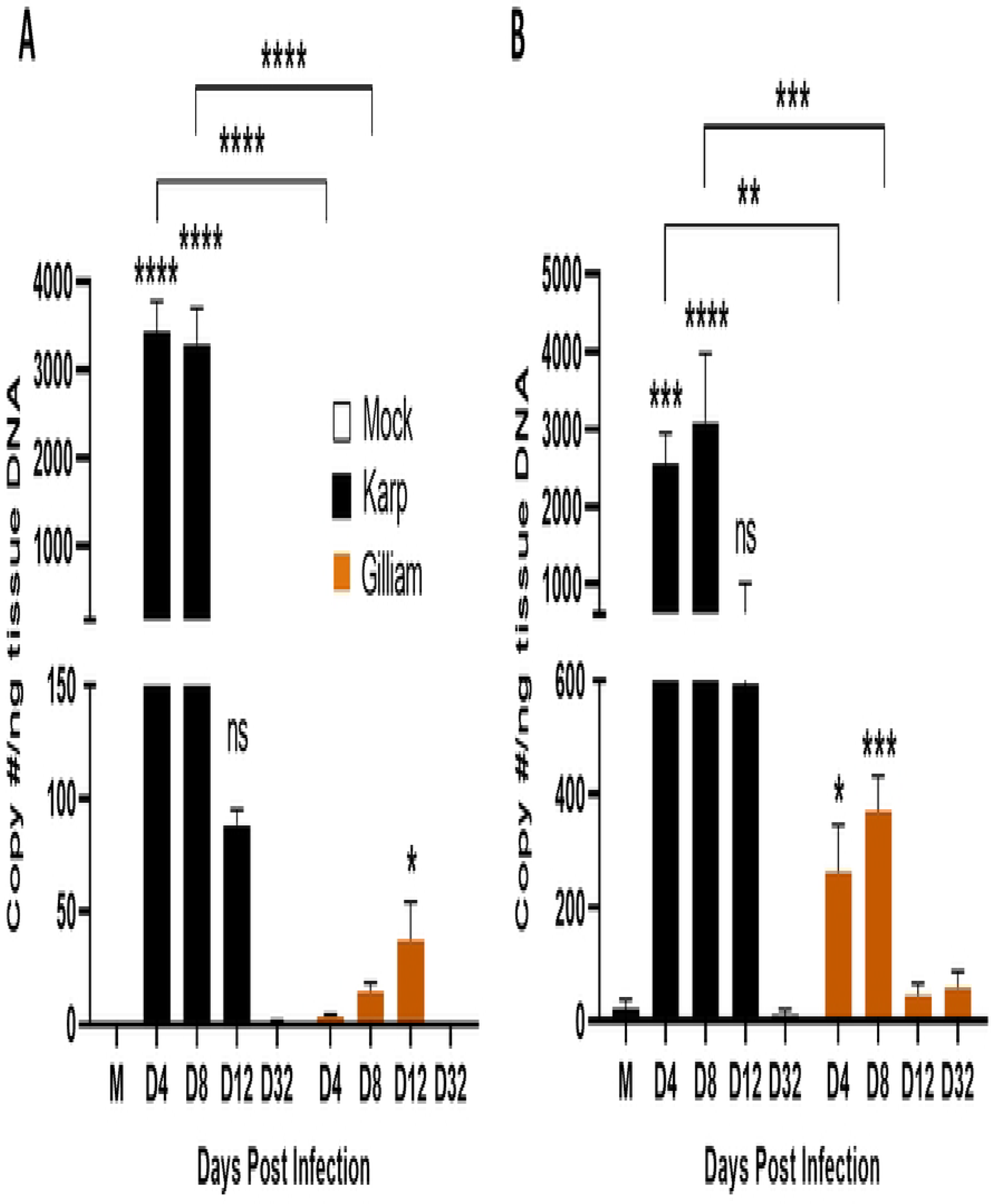
Differential bacterial burdens in *Ot* Karp-vs Gilliam-infected lungs and spleens. C57BL/6 mice were inoculated with Karp, Gilliam, or PBS (mock), as described in Figure 1. A) Lung tissues were collected at days 4, 8, 12 and 32 (5/group); all samples were subjected to DNA extraction and analyzed on the same qPCR plates. B) Spleen tissues were collected similarly (5/group); all samples were subjected to DNA extraction and analyzed. Shown are representative results from two independent studies with similar trends. Data are presented as mean ± SEM. One-way ANOVA (non-parametric) was used for statistical analysis. **, *p* < 0.01; ***, *p* < 0.001; ****, *p* < 0.0001; ns, not significant.

Given that pulmonary infections are common in scrub typhus patients, leading to damage and complications, such as acute respiratory distress syndrome [33], we examined lung pathology during infection with two *Ot* strains. As shown in **Fig 3A and B**, Gilliam-infected lung tissues displayed mild inflammation with limited immune cell infiltration at day 8; the inflammation had lessened by day 12, comparable to the mock levels. In contract, Karp-infected lung tissues showed considerable interstitial pneumonia, as well as pulmonary alveolar edema at day 12 (**Fig 3C**, arrow), indicating an increased vascular permeability. Collectively, our bacteriological and pathological results were consistent with clinical manifestations shown in **Fig 1**, demonstrating that Karp strain was more virulent than Gilliam in mice.

**Fig 3.**
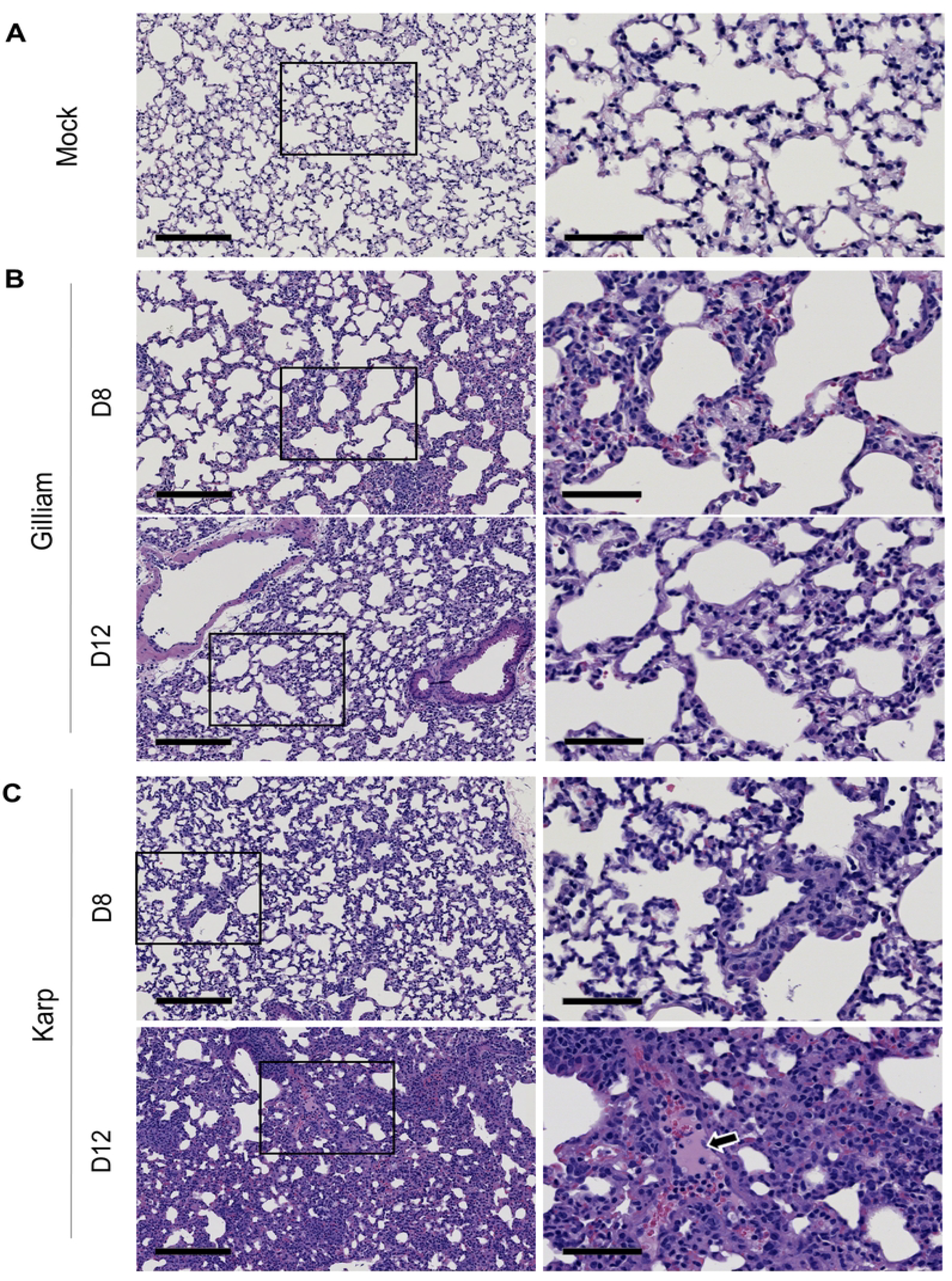
Pathological changes in B6 lungs after infection with *Ot* Karp or Gilliam strain. Mice were inoculated with PBS (mock, A), Gilliam strain (B), or Karp strain (C), as described in Figure 1. Lung tissues (5 mice/group) were collected and subjected to hematoxylin and eosin staining. Pulmonary edema, cellular infiltration, and interstitial pneumonia were observed in Karp-infected tissue at day 12 (white arrow in D12 right). Left scale bars = 200 μm; right scale bars = 70 μm. Shown are representative images from two independent studies with similar results.

### Karp infection results in robust innate and adaptive immune responses in the lungs

Having confirmed the rapid dissemination and growth of Karp strain and severe interstitial pneumonia associated with this infection, we then examined cellular immune responses in the lungs. At indicted days of infection, we perfused lungs with PBS, prepared lung-derived single-cell suspension, and stained cells for flow cytometric analyses. We found that while Karp-and Gilliam-infected lungs showed similar kinetics of immune cell accumulation, which peaked at day 12, Gilliam-infected lungs had approx. 40% lower cell numbers than Karp-infected lungs at day 12 (*p* < 0.0001, **Fig S1**). The analyses of lung innate immune cell subsets and their activation status revealed several unique patterns (**Fig 4A and Fig S2**). 1) While both groups of mice showed nearly identical kinetics of monocyte numbers during days 4-12, Karp-infected lungs had significantly higher numbers of MΦs (CD11b^+^ CD64^+^) and M1-type MΦs (CD80^+^ CD206^-^) at day 12 than Gilliam-infected lungs (*p* < 0.0001). 2) Karp-infected lungs had earlier and sustained influx of neutrophils (PMN) (CD11b^+^ Ly6G^+^), as well as CD63^+^ activated PMNs during days 4-12, than Gilliam-infected lungs. 3) Karp-infected lungs, but not Gilliam-infected lungs, showed progressive accumulation of activated NK cells (CD3^-^ NK1.1^+^ CD44^+^), which peaked at day 12. Both CD4 and CD8 T cells play an important role in *Ot* elimination [8, 34,35]. As shown in **Fig 4B**, Karp-infected lungs showed accumulation of activated CD4^+^ T cells (CD3^+^ CD4^+^ CD44^+^ CD62L^-^), which were approx. 3-fold higher than Gilliam-infected lungs at day 12 (*p* < 0.0001). Likewise, significant accumulation of activated CD8^+^ T cells (CD3^+^ CD8^+^ CD44^+^ CD62L^-^) occurred at day 8 and peaked at day 12, which were significantly higher than Gilliam-infected lungs at day 12 (*p* < 0.0001). Overall, Karp-infected lungs possessed robust and sustained innate immune cell responses, followed by strong CD4^+^ and CD8^+^ T cell responses, all of which consistently peaked at day 12 and were significantly higher than Gilliam-infected lungs. These early robust immune cell expansions in Karp-infected lungs, and to a lesser extent in Gilliam-infected lungs, may play a key role in the over-reactive inflammatory response and lead to more severe clinical manifestations and host mortality.

**Fig 4.**
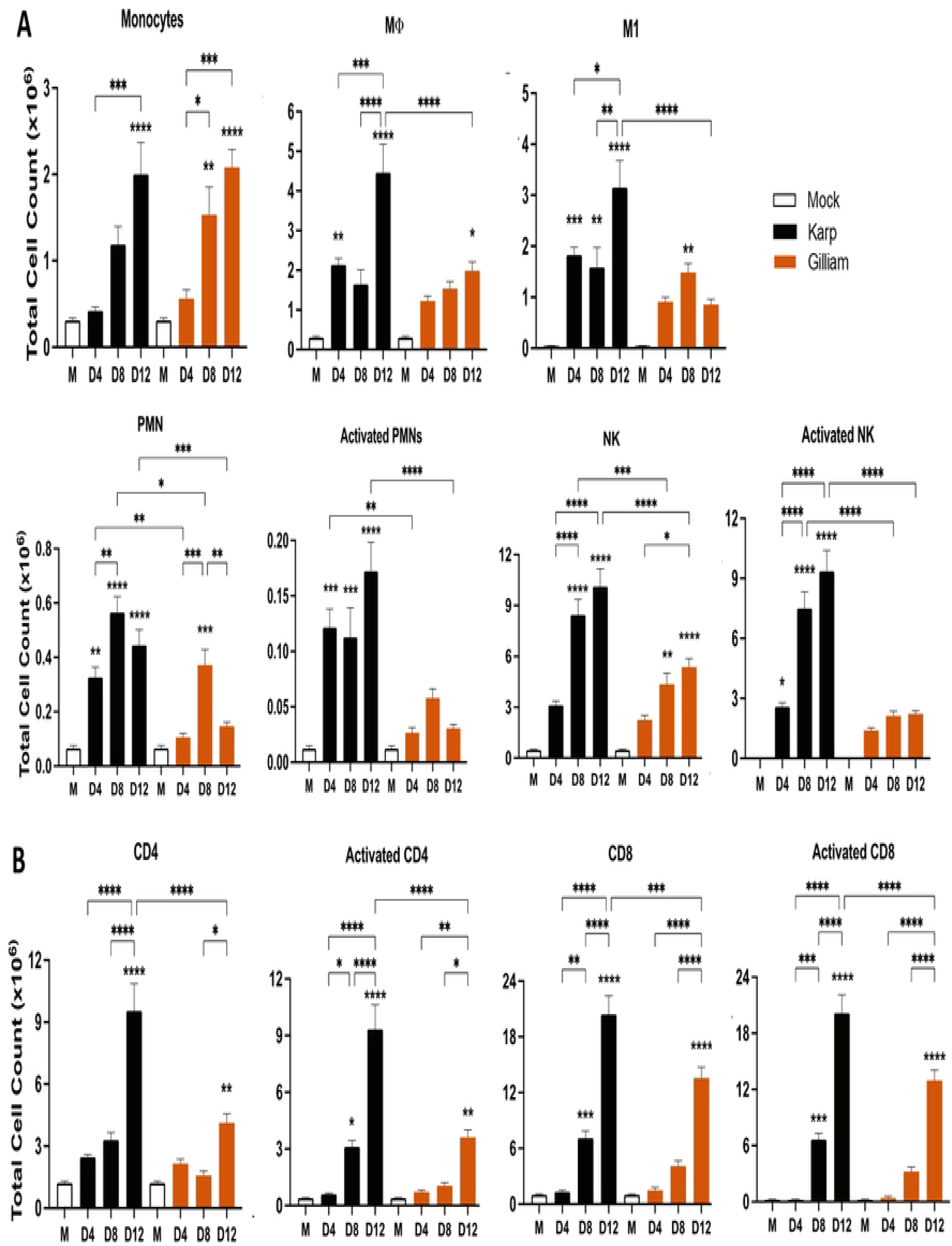
Strong and sustained cellular immune responses in *Ot* Karp-infected lungs. C57BL/6 mice were inoculated with Karp, Gilliam, or PBS (mock), as described in Figure 1. At days 0, 4, 8, and 12 post-infection (5/group), mouse lungs were perfused with a PBS solution. Lung tissues were digested, prepared for single-cell suspension, and stained for flow cytometric analyses (see Materials & Methods for detail). A) Total cell numbers of pulmonary innate immune cell subsets and their activation (in 10^6^). B) Total cell numbers of pulmonary CD4^+^ and CD8^+^ T cells and their activation (in 10^6^). Data are presented as mean ± SEM. Shown are representative results from two independent studies with similar trends. One-way ANOVA was used for statistical analysis. *, *p* < 0.05; **, *p* < 0.01; ***, *p* < 0.001; ****, *p* < 0.0001.

### Karp infection results in high levels of serum cytokines and chemokines

To validate and expand our PCR-based findings, we measured cytokine and chemokine levels in sera by using a Bio-Plex assay. As shown in **Fig 5A**, Karp-infected mice showed statistically significantly elevated levels of CXCL1, CCL2, and G-CSF, especially at day 4; these three markers were significantly increased in Gilliam-infected mice at day 8, but at lower levels than those of Karp groups (*p* < 0.01). By days 8 or 12, Karp-infected mice had significantly higher levels of CCL3 and CCL5 than those of Gilliam-infected mice (**Fig 5B**). Additional markers that were differentially and significantly expressed in one of our independent studies were shown in **Fig S3**, suggesting a trend of higher serum IFN-γ, IL-10, IL-17, and Eotaxin levels in Karp-infected mice around days 8-12. These serum cytokine/chemokine protein profiles were consistent with our findings in the pulmonary flow cytometry results, supporting differential cellular immune response profiles during acute infection with two *Ot* strains.

**Fig 5.**
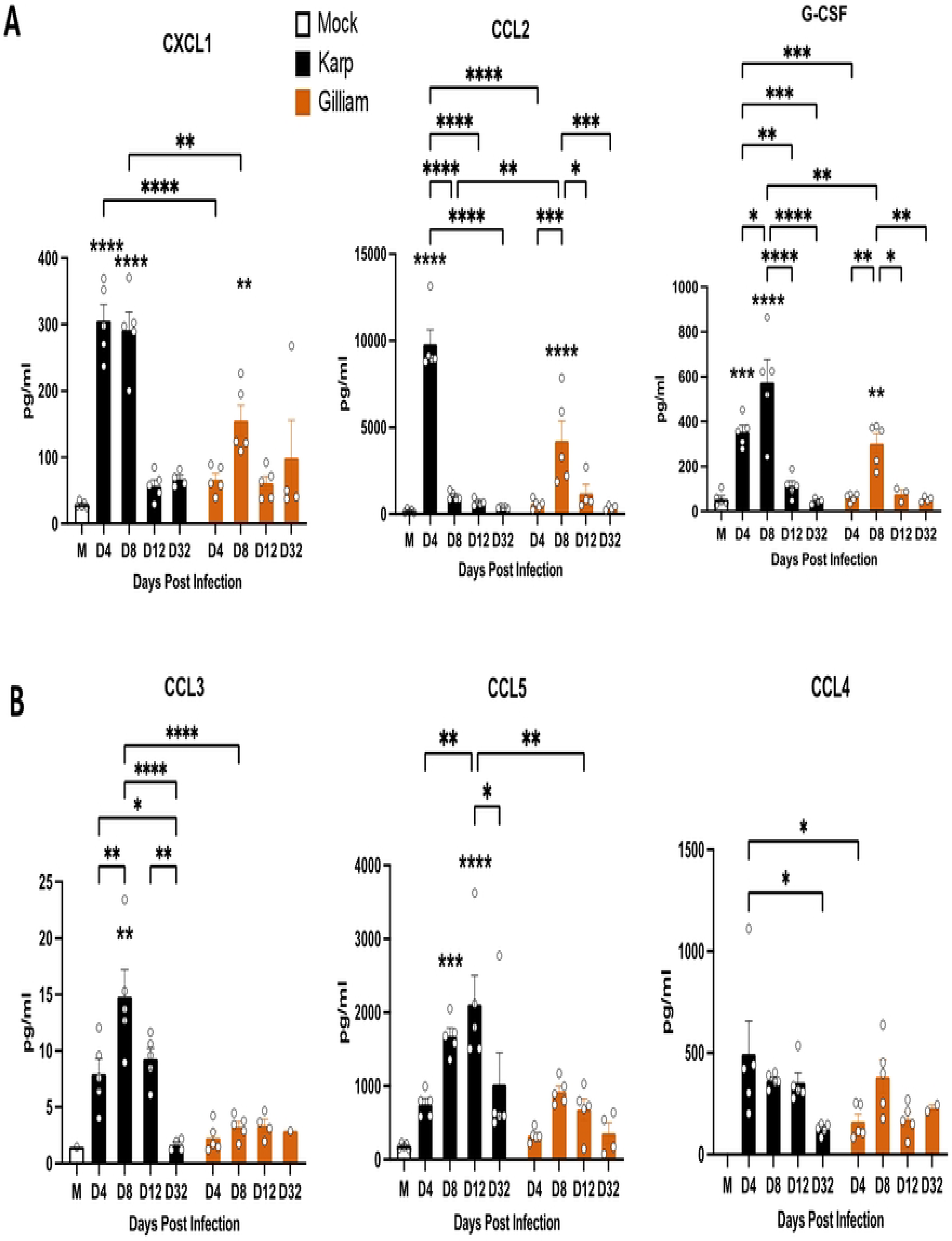
Serum cytokine and chemokine levels in *Ot*-infected mice. C57BL/6 mice were infected, as described in Figure 1. Serum samples were collected at days 4, 8, 12 and 32 and used for cytokine/chemokine measurement via a Bioplex assay (5 mice/group). A) Early biomarkers that were significantly induced at D4 of Karp-, but not Gilliam-infected, mice. B) Significantly and differentially induced markers in Karp-vs. Gilliam-infected mice. Data are presented as mean ± SEM. One-way ANOVA was used for statistical analysis. *, *p* < 0.05; **, *p* < 0.01; ***, *p* < 0.001; ****, *p* < 0.0001.

### Karp infection induces higher lung proinflammatory gene expression as compared to Gilliam infection

To further reveal pulmonary inflammatory response profiles, we examined the gene expression of key cytokine and chemokine genes known to contribute to severe scrub typhus [11]. At day 4, Karp-infected lungs, but not Gilliam-infected lungs, showed significant expression of *CXCL1*, *CXCL2, CCL2*, *CCL3,* and *CCL4*, as compared to the mocks (**Fig 6**). During days 4-12, the *CCL3* and *CCL4* levels were consistently higher in Karp-infected lungs than Gilliam-infected tissues, while both groups showed comparable kinetics and levels of *CCL5.* Of note, Karp-infected lungs showed significantly higher levels of *IFNγ* at days 8 and 12 than Gilliam-infected tissues. Together, these qRT-PCR findings corresponded to the flow cytometry results and serum protein profiles, confirming *Ot* strain-related differential responses in cytokine/chemokine expression.

**Fig 6.**
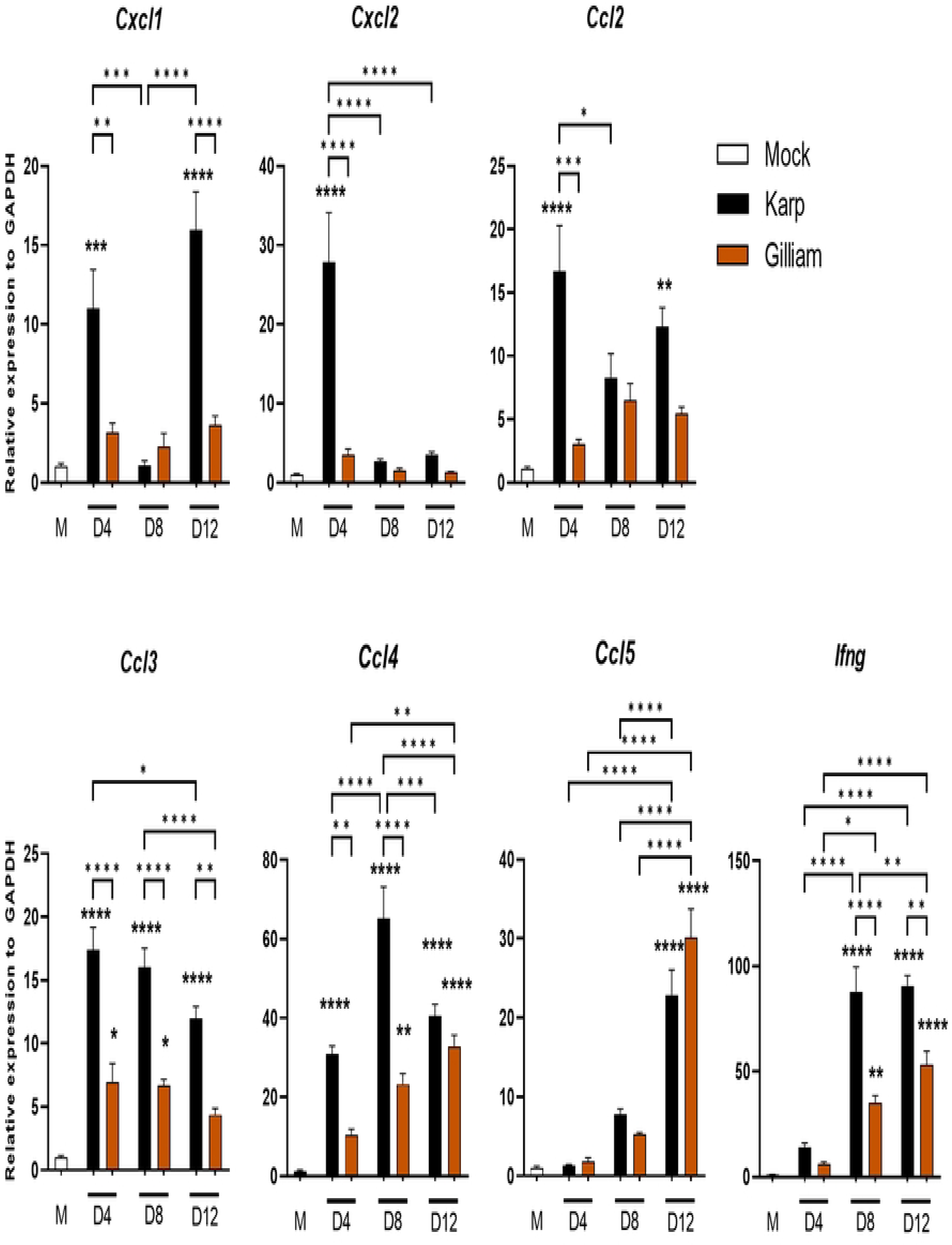
Lung inflammatory gene expression during *Ot* Karp vs. Gilliam infection. C57BL/6 mice were infected, as described in Figure 1. Perfused lung tissues were collected at days 4, 8, and 12 for RNA extraction and qRT-PCR analyses (5 mice/group). Data are presented as values relative to GAPDH and as mean ± SEM. Shown are representative results from two independent studies with similar trends. One-way ANOVA was used for statistical analysis. *, *p* < 0.05; **, *p* < 0.01; ***, *p* < 0.001; ****, *p* < 0.0001.

### Unique MΦ proinflammatory gene profiles are induced by two *Ot* strains

Using the Karp strain, we previously reported that lung MΦ subsets and activation status are strongly associated with disease severity and host mortality [21, 22], and that Karp-MΦ interactions play a key role in orchestrating the innate immune responses, partially via Mincle/FcγR/TNFα-mediated mechanisms [36, 37]. To further investigate differential MΦ responses to different *Ot* strains, we generated bone marrow-derived MΦs from B6 mice and infected cells with Karp or Gilliam (MOI 10) and measured related genes at indicated time points via qRT-PCR (**Fig 7**). At 24 h, Karp infection resulted in a significant upregulation of innate immune genes, including *Il1b*, *Mx2*, *Mincle*, and *Fcgr1*, compared to Gilliam infection (**Fig 7A**). This notable upregulation by Karp was consistently observed at 72 h as well. The proinflammatory cytokines and chemokines *Tnf*, *Cxcl1*, *Ccl2*, and *Ccl5* exhibited significantly higher levels in Karp infection at 72 h (**Fig 7A**), suggesting that the Karp strain induces a hyperinflammatory response in MΦs. Furthermore, Gilliam infection caused higher expression of *Ccl3* and *Ccl4* at 24 h, accompanied by a strong induction of *Egr2* (an M2 marker) at 72 h (**Fig 7B**). Notably, both strains inhibited the expression of *Arg1* at 72 h, which indicates a proinflammatory signature in MΦs infected with *Ot*. Thus, these results are consistent with our findings for *in vivo* study, suggesting the unique activation and polarization of MΦs by diverse *Ot* strains.

**Fig 7.**
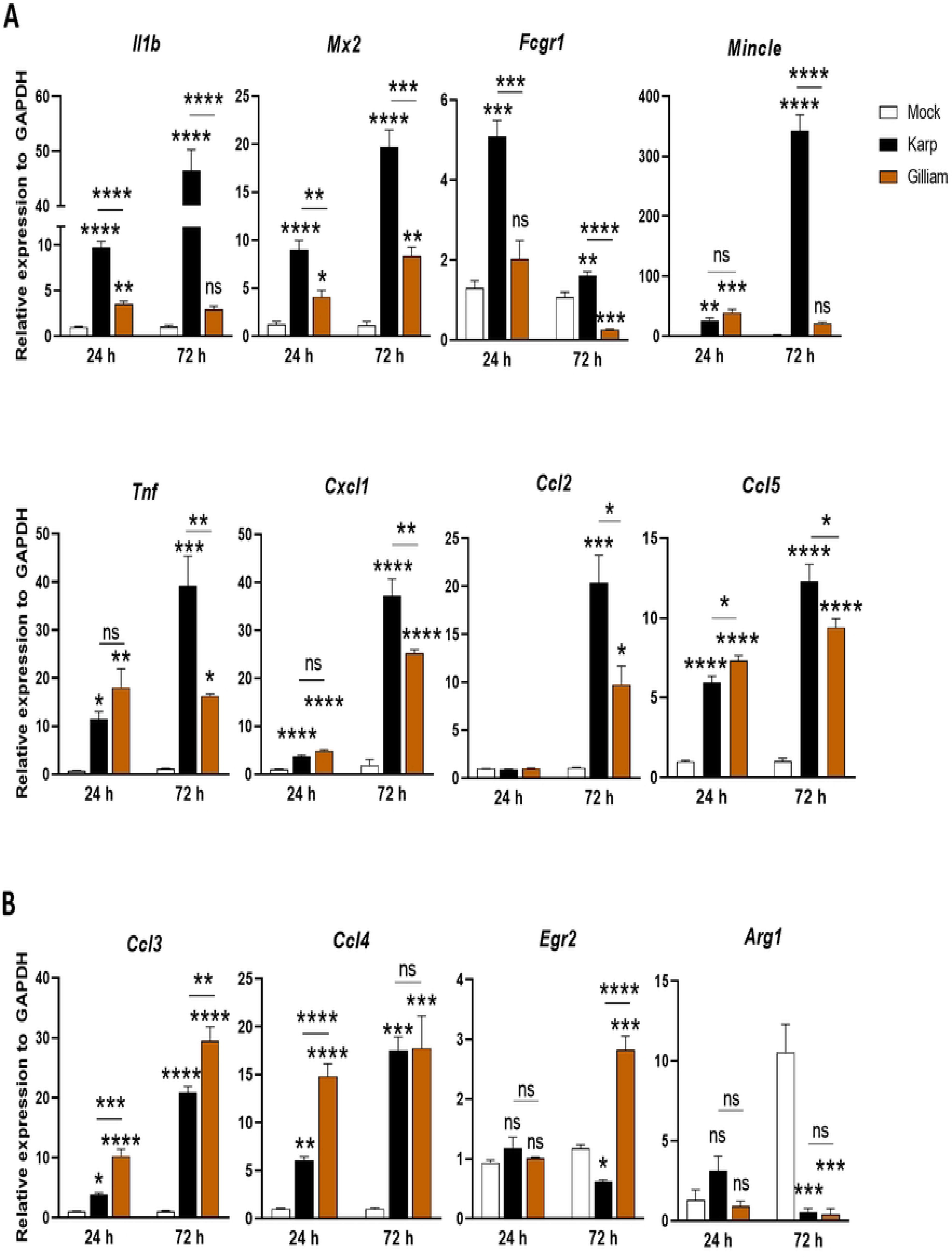
Differential macrophage responses during *Ot* Karp vs. Gilliam infection *in vitro*. Bone marrow-derived MΦs were generated from C57BL/6 mice, seeded in 24-well plates, infected with bacteria (MOI: 10, 4 wells/condition), collected at 24 and 72 h, and analyzed for indicated markers via qRT-PCR. A) Markers preferentially and strongly induced in Karp-infected cells. B) Markers preferentially induced in Gilliam-infected cells, especially at 72 h, or repressed at late stages with both strains. Data are presented as mean ± SEM. One-way ANOVA was used for statistical analysis. *, *p* < 0.05; **, *p* < 0.01; ***, *p* < 0.001; ****, *p* < 0.0001, ns, not significant.

## Discussion

Scrub typhus is a seriously understudied tropical infectious disease, which has recently emerged outside of its traditional geographic areas. Understanding cellular immune responses in the major target organs and during acute stages of infection with different *Ot* strains has been difficult and requires urgent attention. For example, our understanding of the physiologic, antigenic, and virulence differences between Karp and Gilliam is very limited, partially due to the complexity and diversity of *Ot* genome landscapes and the lack of adequate genetic tools for bacterial manipulation [38, 39]. In this study, we conducted the first detailed investigation with these two clinically important *Ot* strains and revealed pulmonary innate and cellular immune responses in the context of distinct clinical outcomes of disease. This study helps understand the potential mechanisms underlying severe scrub typhus vs. self-limited infection and provides several lines of new evidence.

Firstly, we showed murine host-dependent differences in susceptibility to Karp vs. Gilliam infection. While Karp-infected B6 mice showed progressive weight loss and higher disease scores and then reached 20% mortality rates, Gilliam-infected mice gained some weight and showed no signs of disease (**Fig 1A**). Such distinct outcomes of infection were not due to the quality of Gilliam stocks, as an even lower inoculation dose of Gilliam was capable causing lethal outcomes in CD-1 mice (50% mortality rates, **Fig 1B**). These clinical outcomes were consistent with tissue bacterial burdens (**Fig 2**) and lung pathology (**Fig 3**). Corresponding to our recent report [14], we proved here that CD-1 mice were more susceptible to Karp strain than B6 mice, as mortality rates for CD-1 mice reached 100% at day 15 (even though their inoculation dose was lower than that of B6 mice). Therefore, our side-by-side comparison of inbred and outbred mouse models clearly indicated a dose-dependent host susceptibility to two *Ot* strains: Karp-infected CD-1 > Gilliam-infected CD-1 >> Karp-infected B6 >> Gilliam-infected B6. These findings have greatly expanded beyond previous reports for *Ot* strain-based studies [40], offering important murine models for future vaccine-and/or drug-based studies for the control of scrub typhus.

Our conclusion of host-dependent susceptibility is consistent with other murine models for comparison between Karp and Gilliam, which overwhelmingly show Karp is more virulent in the animals, regardless of the routes of infection or mouse strain/stock used [13, 26, 27, 41-45]. For non-human primate models, only a few comparison studies between bacterial strains have been conducted. For example, a recent study with rhesus macaques found that i.d. inoculated Gilliam can cause bigger skin lesions in comparison to Karp inoculation [20]; however, the immunologic or bacteriologic mechanisms underlying such differences have not been explored. The development and improvement of murine scrub typhus models offer great potentials in this regard.

Secondly, we revealed *Ot* strain-related differences in tissue and cellular immune cell responses during Karp vs. Gilliam infection. Our use of perfused pulmonary tissues and multi-color flow cytometry of lung-derived immune cell subsets showed a higher influx and activation of innate immune cell subsets in Karp-over Gilliam-infected mice (**Fig 4, Fig S1**). Pulmonary immune cell comparison represented the most accurate assessment of host immune response at the cellular level, revealing the kinetics and activation status of key cell types during infection. Our most important findings were the following: 1) Karp-infected lungs showed progressive influx/activation of all examined cell types (MΦ, PMN, NK, CD4^+^ and CD8^+^ T cells); nearly all of them peaked at day 12 when lung bacterial burdens were apparently under control (reduced by approx. 40-folds when compared between day 12 vs. day 8, **Fig 2**). Such findings support immune-mediated pathology at severe scrub typhus. 2) As early as day 4, Karp-infected, but not Gilliam-infected, lungs showed significantly increased numbers of MΦ, M1-type MΦ, PMN, activated PMN, and activated NK cell populations, indicating differential host responses at the level of activating myeloid and innate immune cell subsets. Our findings from mouse models were relevant to some observations made from scrub typhus patients. For example, Paris and colleagues have suggested that indicators of PMN activation and neutrophil extracellular trap (NET) formation were considerably increased in severe scrub typhus patients, compared to those with less disease [46]. 3) As early as day 8, Karp-, but not Gilliam-, infected lungs showed significantly increased numbers of activated CD4^+^, total CD8^+^, and activated CD8^+^ T cells. In contrast, Gilliam-infected lungs showed increased activated T cell subsets only at day 12, but to a much less magnitude than Karp-infected lungs. These flow cytometric studies have confirmed and greatly expanded the findings reported in our recent studies with Karp-infected mice [21].

The activation of CD4^+^ T cells in infected mice are consistent with recent study of polyfunctional CD4^+^ T cells in the whole blood of scrub typhus patients [47]. More importantly, our study clearly revealed differential kinetics of cellular immune responses in the lungs during acute stages of infection with two *Ot* strains. Along with NET formation, M1-type MΦs, activated NK, and CD8^+^ T cells can exacerbate tissue damage. The involvement of a proinflammatory state in severe scrub typhus has been noted in other mouse studies [11, 14, 21, 23, 37], as well as in scrub typhus patients[7, 48-50]. We have reported that the stronger innate response in the Karp-infected lungs also promotes excessive T cell activation and responses, which in turn contribute to bacterial control, but also amplify immunopathogenesis [32]. It will be important for future studies to distinguish the monocyte/MΦ functionality between different strain infections to define immune signatures and biomarker relevant to bacterial control and disease outcomes.

Thirdly, we showed differential expression of biomarkers in serum and lung samples during Karp vs. Gilliam infection. We provided evidence that early and high levels of serum CXCL1, CCL2, and G-CSF were hallmarks for Karp-infected mice, which were positively correlated to lung transcript levels of *CXCL1*, *CXCL2*, and *CCL2* at day 4. These chemokines/cytokines might contribute to early and extended recruitment and/or activation of MΦs and PMNs seen in the lungs (**Fig 4**). The importance of some of these proinflammatory factors, including CCL2 and CXCL1, have been established previously and are components of a Mincle-dependent activation response [37]. CCL2 and its receptor, CCR2, have also been suggested to play a role in the influx and activation of monocytes into the lungs, increasing bacterial replication and advancement of interstitial pulmonary inflammation [51]. These levels of host response factors were either markedly mitigated, or showed different kinetics during Gilliam infection, correlating with the greatly reduced bacterial burdens and mild lung pathology. Future studies will be needed to investigate the contributions of specific immune cell subsets and their involvement in the distinct clinical outcomes. Identifying key signatures associated with these immune cell subsets, such as in the study by Luce-Fedrow *et al*. [44], could be useful for parsing strain differences, as well as a future clinical tool to differentiate signatures associated with protective or lethal outcomes [44].

Finally, we revealed differential trends in MΦ’s kinetic responses *in vitro* to help understand host susceptibility to Karp and Gilliam strains (**Fig 7**). We report here the early and significantly higher gene expression of innate responses, such as *Il1b*, *Mx2*, *Mincle,* and *Fcgr1* in Karp-infected MΦs than Gilliam-infected cells. This finding supports our previous report that sensing Karp bacteria via Mincle (a C-type-lectin receptor) and FcγR (Mincle signaling partner) can amplify TNFα production and type 1-skewed proinflammatory immune responses [37]. It is therefore possible that divergent inflammatory signature between Karp and Gilliam infection might be related to Mincle and its downstream signaling [36]. Higher levels of *Tnf*, *Cxcl1* and *Ccl2* in Karp-infected MΦ were consistent with our *in vivo* results (**Fig 6**), showing the increased myeloid cell infiltration and activation in the lungs at early infection. In contrast, Gilliam infection tended to induce higher levels of *Ccl3* and *Ccl4* at either 24 h or 72 h, indicating a unique regulation of inflammatory gene profiles. Although both *Ot* strains repressed *Arg1*, the high levels of *Egr2* (an exclusive M2 MΦ marker [21, 52]) in Gilliam-infected cells suggest a potential of this strain to induce M2-like gene expression, which is in comparison to the Karp strain [21]. It will be of great interest to further examine whether different *Ot* strains can induce unique MΦ activation and polarization profiles via using multi-omics approaches. Such investigation will help define key immune determinants of bacterial clearance, host immune responses, and disease outcomes.

As illustrated in **Fig 8**, our mouse model studies from two *Ot* strains and improved experimental approaches have greatly extended our recent reports for Karp infection in B6 and CD-1 mice, further supporting a notion of type 1-skewed, but type 2-repressed, inflammation in lethal/severe scrub typus [11, 14, 22, 37, 53-55]. The infection by highly virulent Karp strain can lead to uncontrolled bacterial dissemination and replication in the lungs during the early stages. This excessive replication can trigger the production of proinflammatory chemokines (CXLC1/2, CCL2/3, etc.), resulting in the infiltration of innate immune cells (NK, MΦ, neutrophils) and subsequent inflex of activated CD4^+^ and CD8^+^ T cells. These robust proinflammatory responses induced collectively by Karp strain can cause severe damage to pulmonary tissues and result in edema, ultimately leading to the animal’s death during later stages of the infection. In contrast, the less-virulent Gilliam strain can be effectively eliminated at early stages, presumably due to quick and well-regulated activation of innate and adaptive immune cells, leading to limited tissue damage and a subclinical outcome.

**Fig 8.**
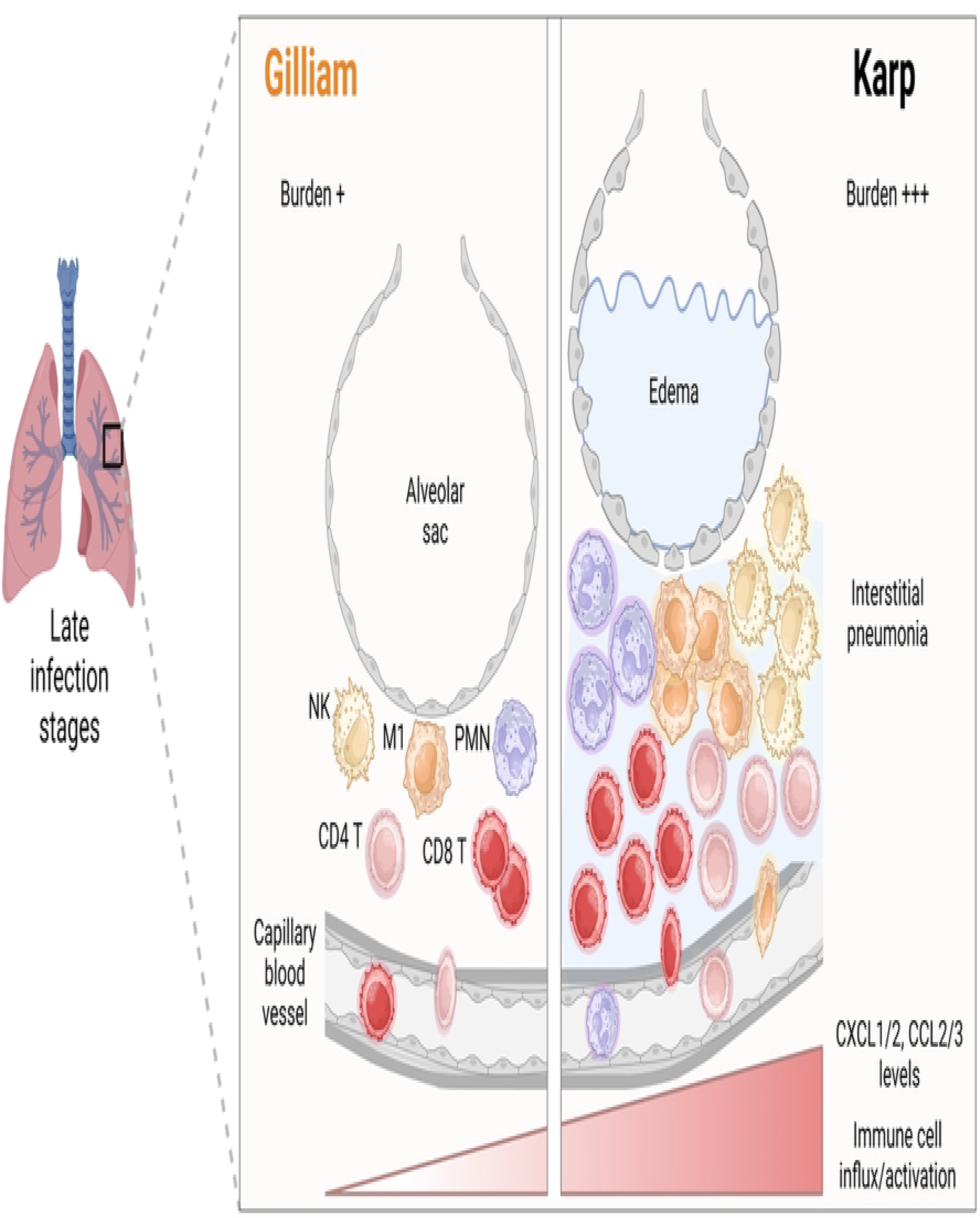
Graphical illustration of differential innate and cellular responses in the lungs during acute infection with *Ot* Karp vs. Gilliam strains. Karp infection causes high tissue bacterial burdens and triggers robust proinflammatory immune responses in the lungs. Strong and uncontrolled production of MΦ and PMN chemotactic factors (CXCL1/2, CCL2/3) and other NK-and T cell-recruiting/stimulating factors, especially at late stages of infection, collectively lead to type 1-skewed cellular responses, acute tissue injury, and host mortality. In contrast, Gilliam dissemination to the lungs and replication in C57BL/6 mice are relatively slow and self-limited, due to mild and balanced cellular responses in the lungs.

At present, the mechanism by which bacteria are eliminated efficiently at the early stage of infection is still unknown. In a recent report for *Rickettsia parkeri*, another closely related obligate intracellular pathogen, it is found that infection control is mediated by both antiviral-like and antibacterial responses, involving IFN-I and IFNγ, respectively [56]. Knowing that IFNγ plays a prominent role in *Ot* control and pathogenesis [57], it will be important to further investigate whether early IFN signaling contributes to the diversity of bacterial clearance in Karp vs. Gilliam infection. Furthermore, it is unclear as to whether different *Ot* strains replicate differently in endothelial cells *in vivo*. Several reports including Mika-Gospodorz *et al.* [58], in which they compared bacterial strains in the context of host response, provide a powerful overview of host-pathogen interaction [58]. Further studies, including the potential use of genetic and molecular tools, will help dissect the underlying mechanism of *Ot* diverse virulence *in vitro* and *in vivo*.

In summary, we have shown that Karp-infected mice produce a higher degree of disease severity, mortality, bacterial burden, and pathologic lesions than that in Gilliam-infected mice. Our findings of strong and uncontrolled activation of cellular immune responses in the lungs, despite of seemingly controlled tissue bacterial burdens at late stages of Karp infection, highlight the contributions of *Ot* strain-dependent, immunopathogenesis in acute tissue injury in scrub typhus. We have provided new evidence for key signaling molecules that are involved in mounting host innate responses and highlighted new research questions and directions. This study provides a framework where strain-specific differences can be analyzed and used for future mechanistic and therapeutic studies.

## Acknowledgements

We would like to thank the UTMB Flow Cytometry and Cell Sorting Core Lab (Meredith Weglarz) for sample analyses, Dr. David Walker for providing the BSL-3 research facilities, and Dr. Ashley Smith for proofreading this manuscript.

## Supporting Information (SI)

**S1 Table. Real-time PCR primers of mouse genes.**

**S1 Fig. Total lung cell counts of infected C57BL/6 mice.** Mice were infected as described in Figure 1. Perfusion was performed on lungs, and corresponding lung lobes were collected at indicated days post-infection. After tissue digestion and cell isolation, the total numbers of lymphocytes were calculated. Data are presented as mean ± SEM. One-way ANOVA was used for statistical analysis. ****, *p* < 0.0001. Shown are representative results from two independent studies.

**S2 Fig. Flow cytometry gating strategy.** A) Lymphocytes were gated according to FSC and SSC, respectively. Live/dead dye-negative cells were identified as live cells. B) Macrophages were gated on CD11b^+^CD64^+^ cells. CD80^+^CD206^-^ macrophages were characterized as M1 macrophages. Monocytes were characterized as CD11b^int^CD64^-^cells.

CD11b^hi^Ly6G^hi^ subpopulation was identified as neutrophils, with CD63 as the neutrophil activation marker. C) Total T cells were gated on CD3 first, followed by CD4 and CD8 gating. CD44^+^CD62L^-^ T cells were considered activated T cells. CD3^-^NK1.1^+^ cells were identified as NK cells, with CD44 as the activation marker.

**S3 Fig. Serum protein levels of cytokine and chemokine in *Ot*-infected B6 mice.** B6 mice were infected, as described in Figure 1. Whole blood was collected for serum preparation at days 4, 8, 12 and 32 and used for cytokine/chemokine measurement via a Bioplex assay (5 mice/group). Protein levels show significant difference between Karp-and Gilliam-infected mice in one of two independent studies. Data are presented as mean ± SEM. One-way ANOVA was used for statistical analysis. *, *p* < 0.05; **, *p* < 0.01; ***, *p* < 0.001; ****, *p* < 0.0001.

